# *Enterococcus faecalis* and *Staphylococcus aureus* mixed species infection attenuates pathogen-specific neutrophil responses and impairs bacterial clearance

**DOI:** 10.1101/2022.05.17.492237

**Authors:** Patrick Hien-Neng Kao, Jun-Hong Ch’ng, Kelvin K.L. Chong, Siu Ling Wong, Kimberly A. Kline

## Abstract

*Enterococcus faecalis* is an opportunistic pathogen that is frequently co-isolated with other microbes in catheterized urinary tract infections and chronically infected wounds. While *E. faecalis* can subvert the host immune response and promote the survival of other microbes via interbacterial synergy, little is known about the impact of immune suppression mediated by *E. faecalis* and how *E. faecalis* impacts the survival of co-infecting microbes. We hypothesized that *E. faecalis* can attenuate neutrophil-mediated responses in mixed-species infection to promote survival of the co-infecting species. Here, we show that *E. faecalis* and *Staphylococcus aureus* mono-species infection activates intracellular ROS production and NET formation, respectively, enabling effective neutrophil-mediated control of the microbial infection. Growth of both bacterial species was enhanced during co-infection in neutrophils *in vitro* and in wounds *in vivo. E. faecalis* reduced *S. aureus*-induced NET formation and *S. aureus* suppressed *E. faecalis-*induced intracellular ROS production. When the species ratios were skewed, the neutrophil reaction profile resembled that elicited by the more abundant species, favoring enhanced survival of the less abundant species. These findings highlight the complexity of the immune response to polymicrobial infections and show that attenuated pathogen-specific immune responses contribute to microbial survival in the mammalian host.

## Introduction

*Enterococcus faecalis*, a common member of the microbiome in the human oral cavity [1], urinary tract [2], and gastrointestinal tract [3-5], has emerged to be one of the leading causes of healthcare-associated infections [6-8]. The ubiquity of *E. faecalis* leads to opportunistic infections in immunocompromised individuals, resulting in cross-transmission among residents in healthcare units [6, 7] or self-contamination when individuals have open wounds that are slow to heal [9]. *E. faecalis* is tolerant or resistant to a variety of antibiotics and also possesses multiple mechanisms to avoid immunosurveillance [10-12], together enabling infections ranging from life-threatening diseases like infective endocarditis and bloodstream infection [13-17] to long lasting infections such as urinary tract infection (particularly catheter-associated UTI (CAUTI)) and wound infection [18-24]. In addition to its association with chronic infections, *E. faecalis* is also often found in polymicrobial infections, with common partners such as *Staphylococcus* species, *Acinetobacter baumannii, Escherichia coli, Klebsiella* species, and *Proteus mirabilis* [18, 25-28] depending on the infection site. Mixed-species infections are often correlated with poorer prognoses such as longer hospital stays, longer intensive care unit stays, higher incidence of septic shock, more in-hospital mortality for bloodstream infections [28-31], failed treatments and higher mortality rates in CAUTI [18], and increased need for amputations for diabetic wound ulcers [32, 33]. Given the poorer outcomes associated with polymicrobial infections, coupled with the fact that antibiotic treatment is often less effective against multi-species communities [34], there is an urgent need to develop alternative therapeutic strategies for common mixed-species infections.

Neutrophils are one of the first responding leukocytes that migrate to injured or infected tissues [35, 36]. Equipped with a wide-array of anti-microbial factors, neutrophils use three mechanisms, to eliminate invading micro-organisms: phagocytosis, degranulation, and NETosis [37, 38]. Historically, neutrophils were considered a type of non-specific immune cell, but more recent findings show that neutrophils react to their microenvironments to provide more specific and efficient responses [39, 40]. For example, signals including different pathogen-associated molecular patterns (PAMPs), cytokines, intercellular interactions, and even surface material can determine the specific responses triggered in neutrophils [41, 42]. Specific and regulated neutrophil responses prevent inappropriate or excessive neutrophil activation which can result in collateral damage to host tissues or precipitate autoimmune reactions [43-45]. Previously, using a mouse model of wound infection, we observed that *E. faecalis* undergo a brief period of acute replication up to one day post infection (dpi) followed by a decline in bacterial load over the next three days, and finally a period of persistent infection that extends until at least seven dpi [46]. Consistent with their role as a first responder to infection [35], neutrophil infiltration is prominent during the first three days of *E. faecalis* wound infection, indicating that these cells play a role in the initial reduction of bacterial burden. However, despite the presence of neutrophils during acute *E. faecalis* wound infection, they are not able to completely clear the infection. Similarly, in a mouse model of *E. faecalis* CAUTI, neutrophils are also recruited within one dpi. While neutrophils reduced *E. faecalis* numbers during CAUTI, neutrophils failed to prevent *E. faecalis* colonization of the catheter [47]. Studies of the interactions between *E. faecalis* and neutrophils have indicated that opsonization, either by complement or antibodies, enable neutrophils to recognize and eliminate *E. faecalis* [48-51]. Characterization of the specific types of neutrophil responses towards *E. faecalis* have not been reported.

*Staphylococcus aureus* is often co-isolated with *E. faecalis*, particularly at wound sites [25]. Most studies have focused on the increased antibiotic resistance that arise when *S. aureus* and *E. faecalis* are grown together. *E. faecalis* can transfer vancomycin-resistant genes to methicillin-resistant *S. aureus* during co-infection [52, 53]; however, little is known what other mechanisms may contribute to the prevalence and persistence of this pair of microbial species. We recently showed that heme cross-feeding from *S. aureus* activates *E. faecalis* aerobic respiration within mature biofilms to increase ATP production and promote growth [54]. While there is an increasing appreciation that most *E. faecalis* infections are polymicrobial in nature, and more studies are establishing the mechanisms by which *E. faecalis* contribute to mixed-species communities [55, 56], investigation into how these *E. faecalis*-containing microbial communities interface with the immune system are limited [57].

In this study, we characterize pathogen-specific responses mediated by neutrophils and evaluate bactericidal effects following co-infection with *E. faecalis* and *S. aureus in vitro* and *in vivo*. Our data demonstrate that neutrophils triggered either intracellular ROS induction or NETosis in response to *E. faecalis* or *S. aureus*, respectively, for optimal bacterial clearance. Notably, *E. faecalis* alone does not induce NETosis, and when it predominates in a mixed-infection, *E. faecalis* can prevent *S. aureus*-induced NETosis to enhance *S. aureus* survival. By contrast, neutrophils fail to augment intracellular ROS levels when *S. aureus* predominated the mixed infection, leading to increased *E. faecalis* CFU. When co-infection occurs at similar numbers, both species experience a moderate benefit of increased survival in the presence of neutrophils. These results show polymicrobial infections can exploit neutrophil-mediated immune responses to promote the overall survival of microbial community.

## Material and Methods

### Bacterial strains and cultures

*Staphylococcus aureus* strain USA 300 was grown in tryptic soy broth (TSB) and *Enterococcus faecalis* OG1RF was grown in brain heart infusion (BHI) medium. To prepare the bacterial inoculum, a single colony was inoculated into 5 ml of liquid broth and grown statically for 16-20 hours at 37°C. *S. aureus* cells were washed and adjusted in PBS to O.D.600 nm= 1.5 (5 × 10^8^ CFU/ml) and *E. faecalis* to O.D.600 nm= 0.5 (3 × 10^8^ CFU/ml) before dilution to the indicated CFU or multiplicity of infection (MOI).

### Colony forming unit (CFU) enumeration

Bacterial suspensions were collected from *in vitro* incubated samples or homogenates of animal wound tissues and serially diluted 10-fold in PBS to achieve 10^7^ -dilution. To measure CFU, dilutions were spotted onto TSB agar (15 %) plates for *S. aureus* and BHI agar plates (15%) for *E. faecalis*. For mixed cultures of *S. aureus* (USA300) and *E. faecalis* (OG1RF), MRSAII selective plate (BIO-RAD, #63757) were used for *S. aureus* and BHI plates containing 25μg/ml of rifampicin were used for *E. faecalis*.

### Mouse neutrophil isolation

Bone marrow neutrophils were freshly isolated from C57BL/6 mice. Tibia and femurs of the hind limbs were collected, and the bone marrow was flushed with 25G needles with ice cold phosphate-buffered saline (PBS). To purify neutrophils from bone marrow cells, magnetic-activated cell sorting (MACS) was performed using a LS column (Miltenyi Biotec, # 130-042-401) and mouse neutrophil isolation kit (Biolegend, #480058) following the manufacturer’s instructions. In short, the cells were suspended in sorting buffer and labeled with biotin-antibody cocktail for 15 minutes on ice. After washing and resuspension, cells were incubated with streptavidin-conjugated nanobeads for another 15 minutes on ice before addition to the LS column installed on a magnetic separator, and the flowthrough was collected as purified neutrophils. In general, 4-6 × 10^6^ neutrophils were collected from one mouse. After isolation, neutrophils were rested in Hank’s balanced salt solution (HBSS) with calcium, magnesium (Thermofisher, #14025076), and additional 10% of mouse serum at 37°C for 30 to 60 minutes before further stimulation or infections.

### *In vitro* neutrophil infection assay

This assay was modified from a previously described study [58]. Briefly, 24-well or 96-well plates (Nunc, ThermoFisher) were coated with 10% fetal bovine serum at 4°C for overnight. The next day bacteria were pre-incubated in HBSS with 10% mouse serum for 30 minutes on ice prior to washing and normalization in HBSS to the required MOI. After resting, neutrophils were seeded into each well in HBSS with 10% of mouse serum (5 × 10^5^ /well for 24-well plate and 1 × 10/well for 96-well plate). Neutrophils were then infected with bacteria in the same volume of HBSS to achieve a final concentration of 5% mouse serum. The infections proceeded at 37°C until the desired timepoint for the as follows: 4-hour incubation for bacterial killing assays and bioimaging assays, 6 hours for ROS detection assay or other specified time points for flow cytometry.

For intracellular ROS assay, neutrophils were pre-incubated with diphenyleneiodonium chloride (DPI, Sigma-Aldrich #D2926-10MG) at 37°C for 30 minutes prior to washing [59]. Cells were then resuspended in Hank’s balance salt solution (HBSS) with 10% of mouse serum before inoculation with bacteria.

### Immunofluorescence microscopy

For immunofluorescence imaging, cells were seeded in either glass-bottom plates (ibidi, μ-plate) or 6-well plates (Nunc, ThermoFisher) with square cover slips inside. At specified times, cells were fixed with 4% paraformaldehyde (Biolegend, #420801) for 10 minutes at room temperature. For staining of citrullinated histone H3, cells were permeabilized with 0.1% triton-x for 10 minutes in PBS. After washing with PBS, the cells were incubated with 2% bovine serum albumin in PBS for 1 hour at room temperature to block nonspecific antibody binding. Samples were then incubated overnight at 4°C with the primary antibody (neutrophil elastase, Abcam #ab68672; citrullinated histone H3, Abcam #ab5103; all were used at 1 : 500 dilution). On the second day, samples were washed and then incubated with secondary antibody for 1 hour at room temperature in the dark. Finally, Hoechst 33342 (100 ng/ml) was used to stain DNA for 15 minutes at room temperature. Images were taken with a Carl Zeiss Axio Observer and cells in each field of view were manually counted. For each sample, at least 10 images were taken and 100-200 cells were counted per sample.

### Extracellular DNA staining and intracellular ROS production

For evaluating extracellular DNA, neutrophils were pre-stained with Hoechst 33342 (100 ng/ml) for 45 minutes in the dark at 37° C after isolation from the bone marrow. The cells were then washed and inoculated with bacteria in the presence of 500 nM of Sytox Orange (ThermoFisher #S11368) to stain extracellular DNA. After 4 hours of incubation at 37°C, cells were washed and fixed for microscopy. One image per sample was taken at 5x magnification and were then analyzed with ImageJ, by which the mean fluorescence intensity of the Sytox Orange signal was calculated.

To detect intracellular ROS production, the DCFDA / H2DCFDA – Cellular ROS Assay Kit (Abcam # ab113851) was used according to the manufacturer’s instructions. Neutrophils were stained with DCFDA for 45 minutes after isolation. Neutrophils were seeded into black-wall glass-bottom 96-well plates (ibidi, μ-plate #89626) and placed in a plate reader (Tecan Infinite^®^ 200 PRO spectrophotometer) immediately after the addition of stimuli. Plates were maintained in the plate reader at 37°C, and Ex/Em of 485/535 signals were recorded for 6 hours at 10 minute intervals. Kinetic data was analyzed by Graphpad Prism 9 and the area under curve was calculated with Microsoft Excel.

### Flow cytometry

Flow cytometer Fortessa X was used to evaluate the surface expression of degranulation markers and integrins. To stain the cells, approximately 8 × 10^5^ neutrophils were collected after the indicated stimulation, washed with PBS, and stained with fixable viability dye (1 : 1000, reconstituted according to manufacturer’s instructions) (ThermoFisher) for 30 minutes at 4°C in the dark. Cells were then washed with 2% FBS in PBS and incubated with Fc blockers (50 ng/ml) (Biolegend, TruStain FcX™ Antibody) for 15 minutes to avoid nonspecific binding. Cocktail of antibodies recognizing CD45 (Biolegend, clone 30-F11), Ly6G (Biolegend, clone 1A8), and CD11b (Biolegend, clone M1/70) were then added in the final dilution of 1:1000 and incubated with cells for 30 minutes at 4°C in the dark. Cells were washed once more with PBS prior to flow cytometry. Analysis of neutrophil viability was performed after gating on neutrophils (CD45+, Ly6G+, CD11b+) with FlowJo, version 10.

### In vivo mouse wound infection

Mouse wound infections were performed as previously described [46]. Briefly, bacteria were grown overnight and normalized in PBS to prepare the inoculum, which was confirmed for accuracy by CFU enumeration. Mice were first anesthetized with ketamine and xylazine. Hair on the dorsal skin was shaved and further removed with hair removal gel. After disinfecting with 70% ethanol, a single wound was created on the dorsal skin by 6-mm biopsy punch. Ten μl of the bacterial inoculum was added to the wound and left to air dry for 3 minutes. A piece of adhesive dressing (Tegaderm™, 3M) was applied to seal the wound. The entire procedure was performed under a heat lamp to prevent hypothermia. 24 hours post infection, a 1 × 1 cm piece of the wound tissue (including the Tegaderm) was collected and homogenized in PBS. Homogenates were then diluted and spotted onto agar plates as described above to determine CFU in the wounds.

### Statistical analysis

GraphPad Prism 9 software was used for statistical analysis. Analytic tests used for evaluating differences among groups with parametric distributions are indicated in each figure legend. Data are presented with mean or mean ± SD of at least 3 biologically independent experiments. P values of < 0.05 were considered statistically significant.

### Ethics statement

Animal procedures were reviewed and approved by Institutional Animal Care and Use Committee (IACUC) in Nanyang Technological University (AUP# A19061) and performed accordingly.

## Results

### Co-infection of *E. faecalis* and *S. aureus* impairs neutrophil-mediated bactericidal effects, favoring survival of the less abundant species in the inoculum *in vitro*

To begin to understand how *E. faecalis* and *S. aureus* polymicrobial infections interface with the immune system, we examined bacterial survival following single and mixed species incubation in the presence of neutrophils *in vitro*. Co-infection of mouse neutrophils with a 1:1 ratio of *E. faecalis*:*S. aureus* resulted in more *E. faecalis* CFU after 4 hours compared to the single-species infection in which *E. faecalis* single-species infection fail to grow to numbers greater than the inoculum **(Fig. 1A)**, and a near doubling of the bacterial CFU in mixed-species infection **(Fig. 1B)**, suggesting that neutrophil-mediated inhibition of *E. faecalis* is impaired in the mixed-species infection. Importantly, all neutrophil infection experiments were conducted in the presence of freshly harvested mouse serum for opsonization required to promote neutrophil-mediated inhibition of *E. faecalis* growth **(Supp Fig. 1)** [60, 61]. *S. aureus* CFU were also increased in the co-infection compared to single-species *S. aureus* infection, albeit to a lesser extent than observed for *E. faecalis* **(Fig. 1A, B)**. While *S. aureus* can promote *E. faecalis* growth in mixed-species mature five day biofilms due to interbacterial metabolic cross feeding [54], augmented *E. faecalis* and *S. aureus* recovery during mixed-species infection of neutrophils was not due to growth synergy in general, because when the two species were co-cultured in the absence of neutrophils for the same duration, the co-culture CFU were not increased **(Supp Fig. 2A, B)**.

**Figure 1.**
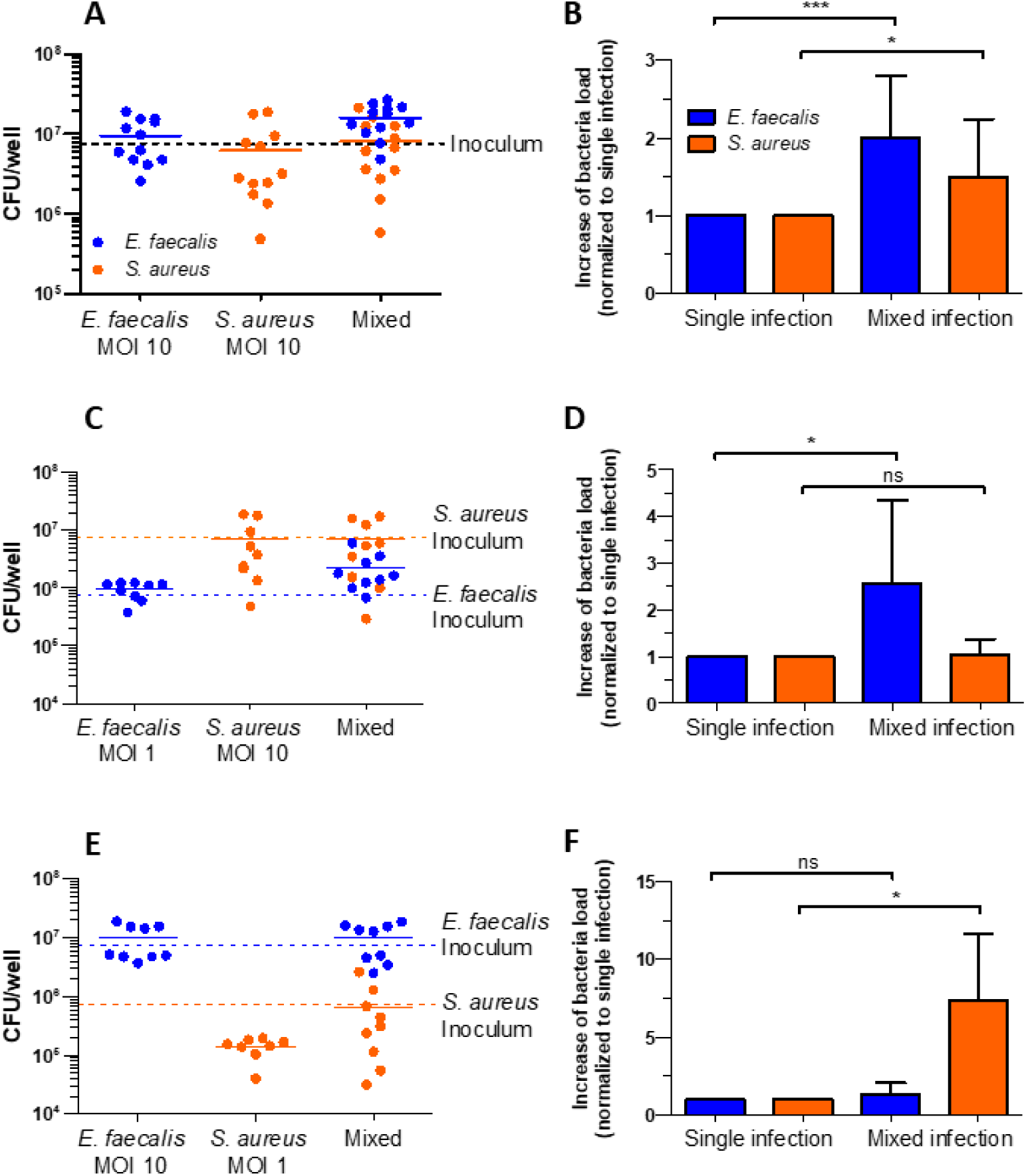
Mixed-species infection results in impaired neutrophil-mediated inhibition, favoring the less abundant inoculating species. CFU following single- and mixed-infection of *E. faecalis* and *S. aureus* after 4-hour incubation with neutrophils were enumerated and normalized. **(A-B)** CFU (A) and fold-change (B) of *E. faecalis* (MOI 10), *S. aureus* (MOI 10), or mixed-infection (MOI 10 + 10). **(C-D)** CFU (C) and fold-change (D) of *E. faecalis* (MOI 1), *S. aureus* (MOI 10), or mixed-infection (MOI *E. faecalis* 1 + MOI *S. aureus* 10). **(E-F)** CFU (E) and fold-change (F) of *E. faecalis* (MOI 10), *S. aureus* (MOI 1), or mixed-infection (MOI *E. faecalis* 10 + MOI *S. aureus* 1). **(A**,**C**,**E)** Horizontal bars represent CFU means and dotted lines to indicate bacterial inoculum. **(B**,**D**,**F)** Bars represent means±SD for fold change relative to each single-species infection. Differences between mixed- and single-species infection were analyzed by paired t-test within each species and differences were considered significant for * p<0.05, *** p<0.001. ns = not significant.

To understand whether one of the two co-infecting species had a dominant impact on the impairment of neutrophil bactericidal activity, we reduced one species in the inoculum by a factor of ten and assessed bacterial CFU post infection. We first infected neutrophils with *E. faecalis* at MOI 1 with *S. aureus* at MOI 10 and observed that only *E. faecalis* CFU significantly increased in the co-infection, whereas *S. aureus* CFU remained comparable between single- and mixed-species infections **(Fig. 1C, D)**. By contrast, when we infected neutrophils with *E. faecalis* at MOI 10 and *S. aureus* at MOI 1, *S. aureus* CFU were significantly increased compared to the single-species infection and *E. faecalis* CFU remained unchanged **(Fig. 1E, F)**. In the absence of neutrophils, co-culture of the two species did not affect *E. faecalis* CFU, regardless of the input inoculum, although *S. aureus* CFU were reduced slightly **(Supp Fig. 2C-F)**. These results suggest that co-infection protects the less abundant species from neutrophil-mediated inhibition, thus favoring its survival.

### Co-infection of *E. faecalis* and *S. aureus* increases bacterial CFU *in vivo* during wound infection, favoring survival of the less abundant species in the inoculum

To determine whether the enhanced survival of bacteria in mixed-species infection observed *in vitro* also occurs in vivo, we used a mouse wound infection model, introducing *E. faecalis* and *S. aureus* at different inoculum ratios and enumerating CFU in the wound tissue 24 hours post infection (hpi). This model was chosen because strong infiltration of neutrophil was observed within 24 hpi, providing a reasonable surrogate for the *in vitro* infection experiments [46]. Following inoculation of the wound with a 1:1 ratio of *E. faecalis* and *S. aureus* (10^6^ CFU of each), similar to the *in vitro* experiments, *E. faecalis* CFU were approximately 7-fold greater compared to the single-species infection **(Fig. 2A)**. By contrast, *S. aureus* replicated to higher numbers than *E. faecalis* and CFU were not enhanced in the mixed-species infection **(Fig. 2A)**. To determine whether reducing *S. aureus* in the input ratio would favor *S. aureus* growth in vivo, as we observed for 1:10 *E. faecalis*:*S. aureus* co-infection *in vitro*, we inoculated wounds with a reduced *S. aureus* inoculum while keeping the *E. faecalis* inoculum at 10^6^ CFU. This mixed-ratio infection resulted in significantly greater CFU for both microbes compared to single-species infection, with *E. faecalis* CFU increased slightly (less than 3-fold), and *S. aureus* increased by more than 100-fold **(Fig. 2B)**. These results correlated with *in vitro* observations in which the sub-dominant species at the time of inoculation was better protected from neutrophil killing during co-infection.

**Figure 2.**
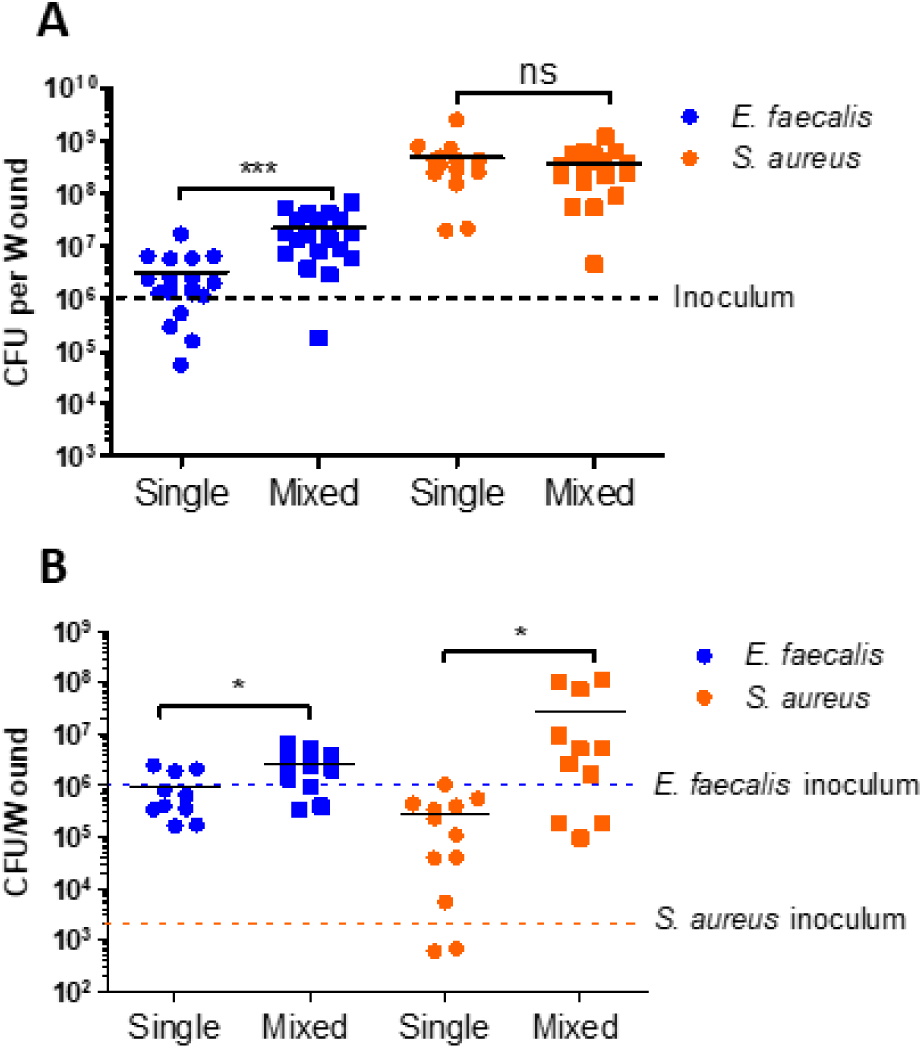
Co-infection promotes bacterial survival and replication in mouse wounds, favoring the less abundant inoculating species. CFU of bacteria recovered from wounds 24 hours post infection. **(A)** Mouse wounds were infected with *E. faecalis* (10^6^), *S. aureus* (10^6^), or both (10^6^ of each). **(B)** Mouse wounds were infected with *E. faecalis* (10^6^), *S. aureus* (2 × 10^6^), or both. Horizontal bars represent CFU means and dotted lines to indicate bacterial inoculum. Differences between mixed- and single-species infection were analyzed by unpaired t-test within each species and differences were considered significant for * p<0.05, *** p<0.001. ns = not significant.

### *E. faecalis* induces intracellular ROS production in neutrophils, which is reduced by *S. aureus*

Opsonized *E. faecalis* can be phagocytosed by neutrophils followed by intracellular ROS induction; however, the level of ROS production does not necessarily correlate with neutrophil-mediated killing [51, 62]. To first evaluate the role of ROS in the inhibition of *E. faecalis* in our infection model, we pretreated neutrophils with diphenyleneiodonium chloride (DPI) to inhibit intracellular ROS production [59] before infection with *E. faecalis*. Compared to an approximately 80% reduction in *E. faecalis* CFU by vehicle-treated neutrophils, ROS inhibition resulted in only a 50% reduction in CFU **(Fig. 3A, B)**. DPI treatment was not toxic to neutrophils **(Supp Fig. 3)**, showing that impaired *E. faecalis* inhibition by DPI-treated neutrophils was not a result of reduced neutrophil viability, together demonstrating that intracellular ROS contributes to neutrophil-mediated inhibition of *E. faecalis*. We next measured intracellular ROS production after single- and mixed-species infection at a 1:1 ratio (MOI 10 of each species) by pre-staining neutrophils with DCFDA before washing and infection, such that only ROS produced within neutrophils can be detected. *E. faecalis* induced intracellular ROS production in neutrophils at similar levels to the TBHP positive control **(Fig. 3C)**. By contrast, not only did *S. aureus* fail to stimulate detectable ROS production following single-species infection, it also significantly reduced the *E. faecalis*-induced ROS during co-infection to an intermediate level between *E. faecalis* and *S. aureus* single-species infection **(Fig. 3C)**. Lowering the *E. faecalis* inoculum to MOI 1 resulted in delayed and lower ROS levels, which was still significantly decreased in the presence of *S. aureus* MOI 10 **(Fig. 3D)**. By contrast, when *S. aureus* was outnumbered by *E. faecalis* 1:10, ROS production in co-infection was comparable to the *E. faecalis* single-species infection **(Fig. 3E)**. These results show that neutrophils respond to *E. faecalis* with intracellular ROS production, which can be reduced in a *S. aureus*-dose-dependent manner. Together these data suggest that *S. aureus*-mediated reduction of ROS, which is normally induced by *E. faecalis* and contributes to its inhibition, can promote *E. faecalis* survival during co-infection.

**Figure 3.**
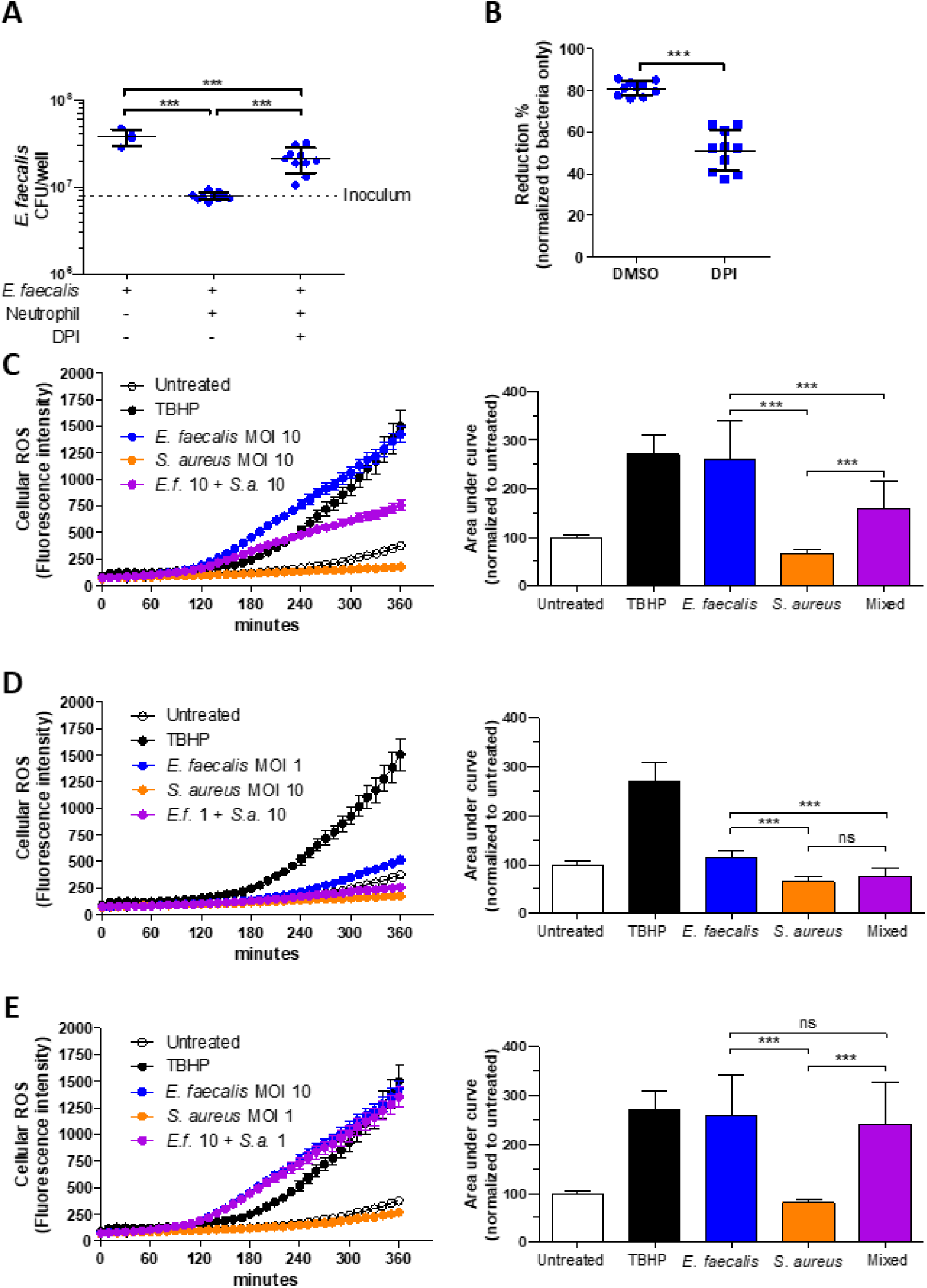
Intracellular ROS in neutrophils is responsible for *E. faecalis* inhibition and is reduced by *S. aureus* during co-infection. **(A-B)** Neutrophils were pretreated with vehicle (DMSO) or DPI (30 μM) for 30 minutes at 37°C before infection with *E. faecalis. E. faecalis* CFU were calculated (A) and percent reduction compared to bacterial alone were enumerated (B). All data are presented with means±SD and dotted line on the CFU graph indicates *E. faecalis* input. **(C-E)** Neutrophils were prestained with DCFDA for 30 minutes at 37°C prior to washing and infection. ROS produced by neutrophils upon exposure to bacteria during the following 6 hours was measured, with left panels showing ROS kinetics and right panels showing area under the curve as a measure of overall ROS production. Various inoculates were used, with *E. faecalis* MOI 10 plus *S. aureus* MOI 10 (A, B), *E. faecalis* MOI 1 plus *S. aureus* MOI 10 (C, D), and *E. faecalis* MOI 10 plus *S. aureus* MOI 1 (E, F). Statistics was analyzed with either one-way ANOVA (A, C-E) or unpaired student t-test (B) and differences were considered significant for *** p<0.001. ns = not significant.

### *E. faecalis* co-infection reduces NETosis induced by *S. aureus*

While *S. aureus*-mediated reduction of intracellular ROS correlates with increased *E. faecalis* protection from neutrophil killing in co-infections, the question remained of how co-infection similarly protects *S. aureus* from neutrophil killing. Since stimulation of neutrophil NETosis contributes to the clearance of *S. aureus* [58], we hypothesized that co-infection with *E. faecalis* may interfere with the formation of NETs. However, the ability of *E. faecalis* to induce NETosis had not been previously reported. Therefore, we examined three markers of NETosis following *E. faecalis* single-species infection and found no evidence for decondensed chromatin, extracellular DNA (as measured by cell-impermeant Sytox Orange), or extruded neutrophil elastase **(Fig. 4A-E)**. By contrast and as demonstrated previously by others [58], *S. aureus* infection resulted in significant induction of all three markers, each of which were significantly reduced following a 1:1 co-infection with *E. faecalis* **(Fig. 4A-E)**, indicating that the presence of *E. faecalis* interferes with *S. aureus*-induced NET formation. Given that the NETosis markers may also be observed when neutrophils lose membrane integrity or upon other forms of cell death, we further evaluated the level of citrullinated histone H3 as a measure of bona fide NETosis. Histone citrullination, which contributes to chromatin decondensation observed during NET formation, is an active response carried out by peptidylarginine deiminase 4 (PAD4) upon various stimulation, including bacterial infection [63, 64]. Consistent with other NETosis markers, *E. faecalis* single-species infection resulted in very few neutrophils containing citrullinated histone H3, similar to uninfected controls **(Fig. 4F, Supp Fig. 4)**. By contrast, *S. aureus* infection resulted in approximately 40% of total neutrophils with citrullinated histone H3, which was partially yet significantly reduced in mixed-species infection **(Fig. 4F, Supp Fig. 4)**. Upon closer inspection, the signals of citrullinated histone H3 (red) were in proximity with *S. aureus* (green), and primarily associated with cell aggregates **(Supp Fig. 5)**. However, in mixed-species infection, *S. aureus*-containing neutrophils were negative of citrullinated histone H3 (white arrowhead) **(Supp Fig. 5)**, suggesting histone citrullination is the pathway suppressed by *E. faecalis* during mixed-infection, leading to attenuated NETosis. *E. faecalis*-mediated NETosis reduction was lost when its ratio in the inoculum was reduced to MOI 1 **(Fig. 4G)**. Reducing the *S. aureus* MOI to 1 resulted in fewer neutrophils undergoing NETosis as assessed by decondensed chromatin, as compared to MOI 10 (∼50% to 30%, respectively), and this number was significantly reduced during co-infection with *E. faecalis*, to levels comparable to *E. faecalis* single-species infection **(Fig. 4H)**. These data show that *E. faecalis* interferes with *S. aureus*-induced NETosis during co-infection, which explains why co-infection with *E. faecalis* results in increased *S. aureus* survival.

**Figure 4.**
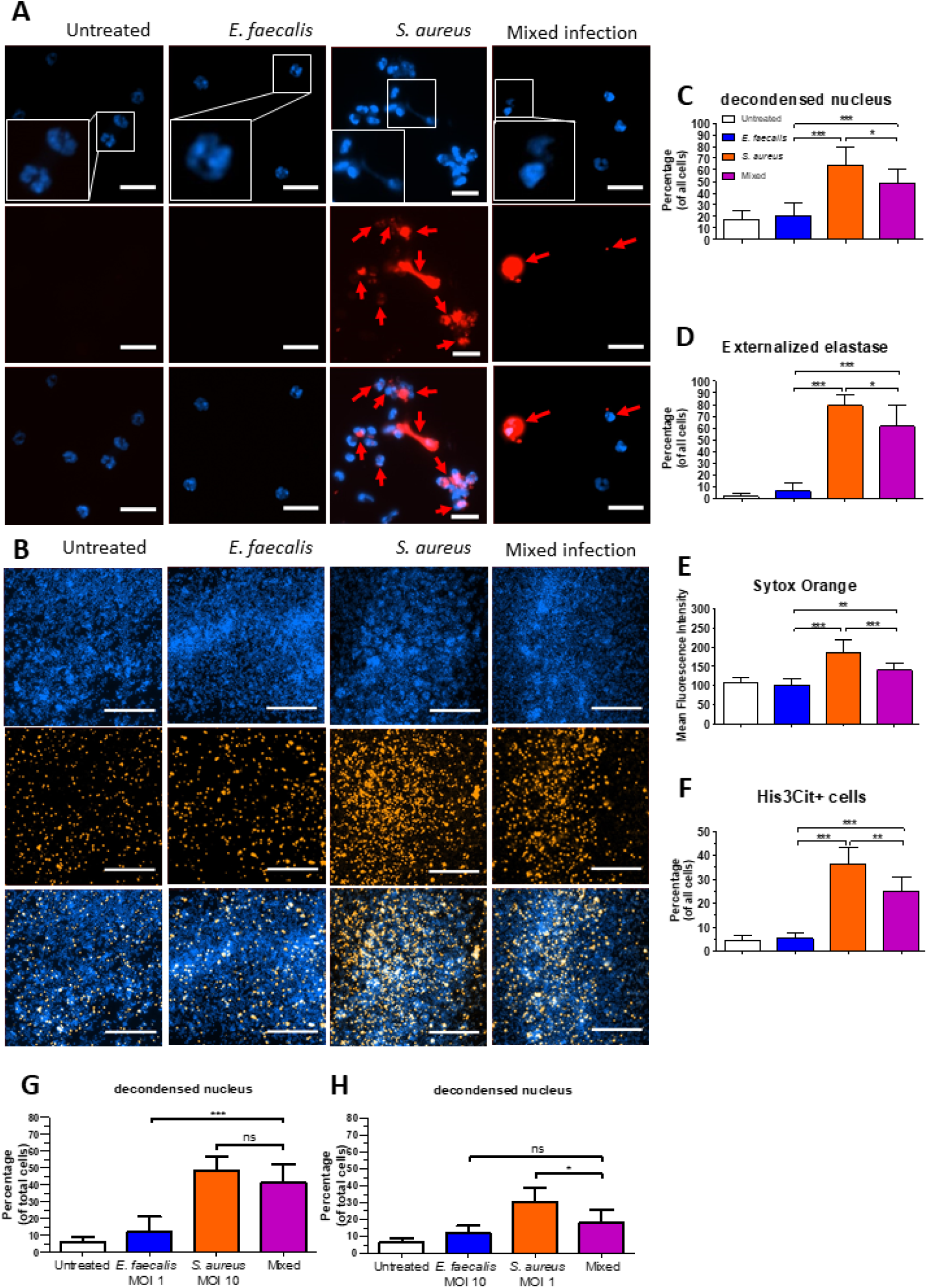
*E. faecalis* reduces *S. aureus*-induced NETosis. Evaluation of NETosis following single- and mixed-infection after 4-hour incubation. **(A-E)** Neutrophils infected with MOI 10 of *E. faecalis* and/or *S. aureus*. **(A)** NETosis was detected by examining the nuclear morphology and externalized neutrophil elastase. Representative images of neutrophil DNA stained with Hoechst (blue) and immunofluorescence for neutrophil elastase (red) (scale bar = 20μm). **(B)** Extracellular DNA was measured as a surrogate marker for NETosis. Neutrophils were pre-stained with Hoechst to label DNA (blue) and after 4-hour incubation the extracellular DNA was stained by Sytox Orange, a non-membrane-permeable DNA dye (scale bar = 200μm). For quantification, 3 independent experiments were performed. The percentage of **(C)** cells with decondensed chromatin, **(D)** cells expressing extruded neutrophil elastase, and **(F)** cells with citrullinated histone H3 were evaluated by examining over 100 cells per experiment. **(E)** Mean fluorescence intensity of Sytox Orange for eDNA was measure and analyzed by ImageJ. **(G-H)** Morphology of neutrophil nucleus was examined to evaluate the level of NETosis after 4-hour incubation with different inoculum of *E. faecalis* and/or *S. aureus*. **(G)** MOI 1 of *E. faecalis* and MOI 10 of *S. aureus* were used for infection. **(H)** MOI 10 of *E. faecalis* and MOI 1 of *S. aureus* were used for infection. All quantified results are presented with means±SD. Statistics was analyzed with one-way ANOVA and differences were considered significant for * p<0.05, ** p<0.01, *** p<0.001. ns = not significant.

## Discussion

Polymicrobial infections, often correlated with chronic infections such as cystic fibrosis and diabetic ulcers, are serious medical challenges and complicated treatment regimens raise the risk of antibiotic resistance. Microbial communities use various mechanisms to promote their survival, including metabolite exchange as energy sources for other microbes [54, 56, 65], establishing biofilms which promote persistence in the face of immune and antibiotic activities [66, 67], and induction or transfer of virulence factors among members [52, 68]. While host-microbial community interactions have been investigated, existing studies have focused on the immunomodulating effect of one species on the outcome of the coinfecting species’ infection [57]. Here we demonstrate that co-infection impacts the immune response by diverting or suppressing pathogen-specific responses, which leads to overall impaired microbial clearance. This study identified specific, primary murine neutrophil reactions toward *E. faecalis* and *S. aureus*, namely intracellular ROS production and NETosis, respectively. During mixed-ratio co-infection, the neutrophil response phenotype favored that which occurs in response to the more abundant species, which coincides with attenuation of the bactericidal effects against the minor species. A similar outcome was also observed in a mouse wound infection model, where the abundant species derive a benefit in the mixed-infection, indicating the impaired neutrophil response may also take place *in vivo*. In addition to neutrophil-specific responses, other host cells (e.g., platelets, fibroblasts, and epidermal cells) or bacterial metabolite exchange may also contribute to the enhanced colonization in the *in vivo* environment, and further investigation is underway to understand the spectrum of interactions that occur between the polymicrobial community and the host.

While opsonization is required for efficient neutrophil killing of *E. faecalis* [61, 69], the mechanism by which neutrophils kill this species has not been reported. We show that *E. faecalis* stimulates neutrophils to produce intracellular ROS that leads to *E. faecalis* clearance. ROS is an important regulator of NET formation [70]; however, *E. faecalis*-induced ROS does not lead to NETosis. This is not unprecedented, given that neutrophils stimulated with GM-CSF and LPS also produce intracellular ROS that does not coincide with NET formation [71]. One potential explanation is that *E. faecalis*-induced ROS accumulates within phagosome which, together with fusion of primary granules to the phagosome [72, 73], could result in sequestration of NETosis activators such as neutrophil elastase (NE) and myeloperoxidase (MPO) [74], eventually leading to the inhibition of *S. aureus*-induced NETosis in mixed-infection. However, it is unclear whether the reduction of histone citrullination also results from the sequestration of NET-inducers like NE and MPO, or whether *E. faecalis* possesses other mechanisms to directly reduce the activity of PAD4.

While *E. faecalis* alone fails to induce NETosis, and co-infection of *E. faecalis* reduces the levels of *S. aureus*-mediated NET production, a significant amount of NETosis still occurs in mixed-species infections, especially when *S. aureus* is in greater abundance. However, *E. faecalis* does not appear to be susceptible to NETosis-mediated killing given that *E. faecalis* proliferates during mix-species infections. Microbial virulence factors such as capsule and endonuclease help pathogens avoid NET killing activity or degrade NET structure [45, 75]. The *E. faecalis* strain used in these studies (OG1RF) does not encode capsule or any predicted nucleases, so it remains to be determined how *E. faecalis* escapes NET-mediated inhibition.

Conversely, we also demonstrate that co-infection with *S. aureus* reduces *E. faecalis*-induced intracellular ROS production, which correlates with impaired inhibition of *E. faecalis*. Several *S. aureus*-derived factors such as SaeRS-regulated factors and lipoic acids have shown suppressive effects towards ROS production in neutrophils and macrophages [76, 77]. In addition, NETosis may also interfere with intracellular ROS accumulation by inhibiting phagocytosis and shift the balance toward extracellular ROS release [78, 79]. Although beyond the scope of this study, the mechanism by which *S. aureus* affects the reduction of *E. faecalis*-driven intracellular ROS accumulation should be addressed. Surprisingly, in our experiments *S. aureus* infection does not induce detectable intracellular ROS production. In previous reports, *S. aureus* infection induces transient intracellular ROS production followed by prolonged hypochlorite production, as well as sustained extracellular hydroperoxide production [77]. Hypochlorite is derived from hydroperoxide by MPO and is crucial for NET formation [80, 81]. It is possible that *S. aureus* infection activates neutrophils to rapidly transform intracellular hydroperoxide into hypochlorite, which is not detectable by the DCFDA/H2DCFDA assay. This may suggest that *E. faecalis* and *S. aureus* activate neutrophils to produce different types of ROS. In addition to suppressing ROS production, *S. aureus* also tolerate ROS-mediated killing via factors such as methionine sulfoxide reductases [82], which may explain how *S. aureus* survive in an ROS-producing mixed-species infection.

In conclusion, we have demonstrated that neutrophils undergo distinct responses toward *E. faecalis* and *S. aureus* to achieve effective bacterial clearance with skewed responses toward the dominant species in a multi-species infection leading to impaired microbial inhibition. Immunotherapies of the future that are designed to promote immune clearance of pathogens must factor in this complexity given that infections are often polymicrobial in nature.

Abbreviations: Definition
ATP: Adenosine triphosphate
BHI: Brain-heart infusion
CAUTI: Catheter-associated urinary tract infection
CFU: Colony-forming unit
DCFDA: Dichlorofluorescin diacetate
dpi: Days post infection
DPI: Diphenyleneiodonium
DMSO: Dimethyl sulfoxide
*E.f.*: *Enterococcus faecalis*
H3Cit: Citrullinated histone H3
HBSS: Hanks’ balanced salt solution
hpi: Hours post infection
MACS: Magnetic-activated cell sorting
MOI: Multiplicity of infection
MPO: Myeloperoxidase
NE: Neutrophil elastase
NET: Neutrophil extracellular trap
NMS: Normal mouse serum
NS: Not significant
PAD4: Peptidylarginine deiminase 4
PAMP: Pathogen-associated molecular pattern
PBS: Phosphate buffered saline
ROS: Reactive oxygen species
*S.a.*: *Staphylococcus aureus*
TBHP: tert-butyl hydrogen peroxide
TSB: Tryptic soy broth

---

**Supp Figure 1.**
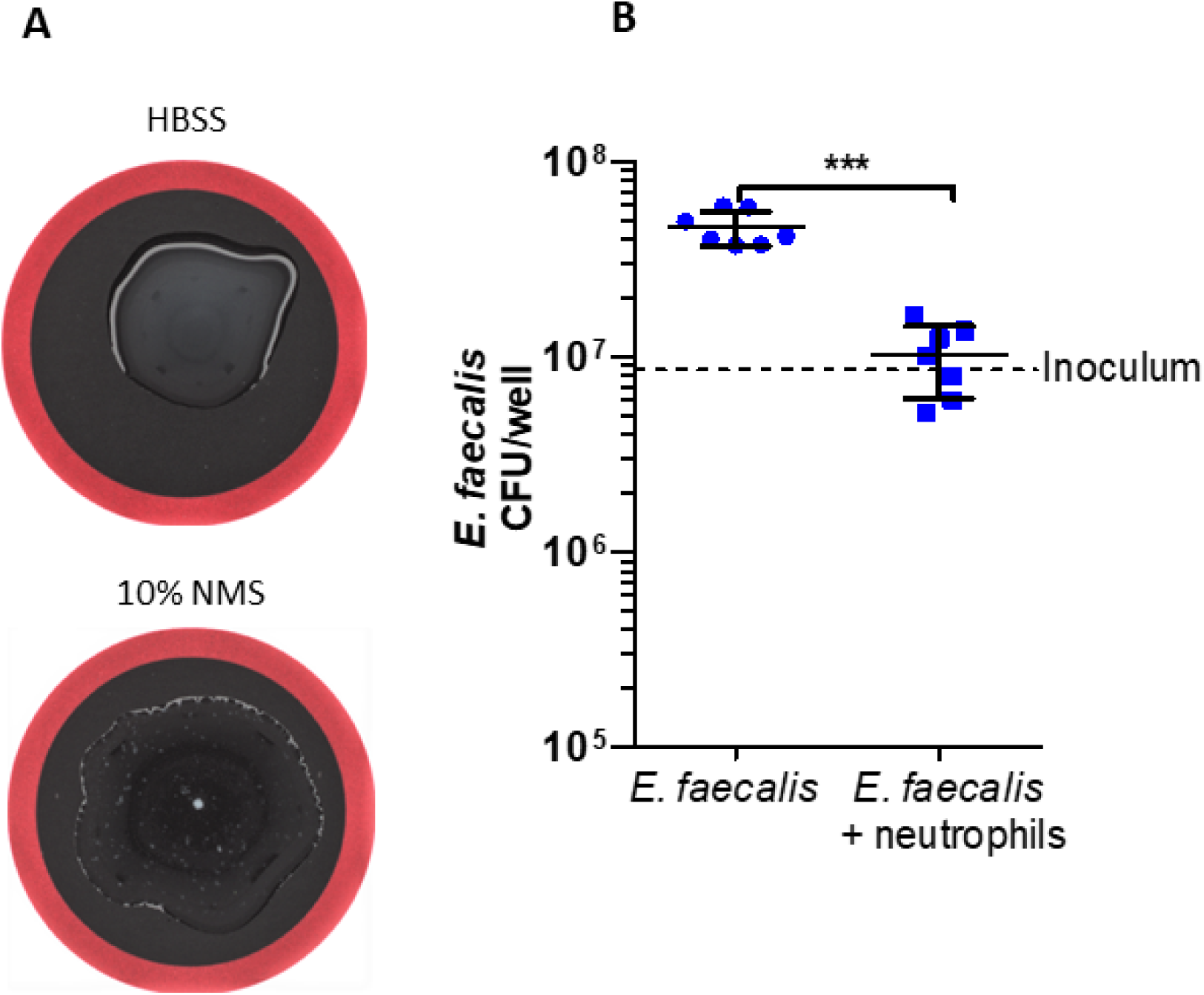
Serum opsonization *E. faecalis* promotes neutrophil inhibition of *E. faecalis*. **(A)** Agglutination of *E. faecalis*. HBSS and HBSS supplemented with 10% normal mouse serum were used to incubate *E. faecalis* in the absence of neutrophils for 30 minutes at room temperature to observe aggregation (particles) in bacterial suspensions. **(B)** CFU of *E. faecalis* recovered after 4-hour incubation with or without neutrophils. 0.1% of Triton-x was added to lyse neutrophils and release intracellular *E. faecalis*. **(B)** Bars represent means±SD for CFU and dotted lines to indicate bacterial inoculum. Differences between *E. faecalis* only group and *E. faecalis* with neutrophils group were analyzed by unpaired t-test within each species and differences were considered significant for *** p<0.001.

**Supp Figure 2.**
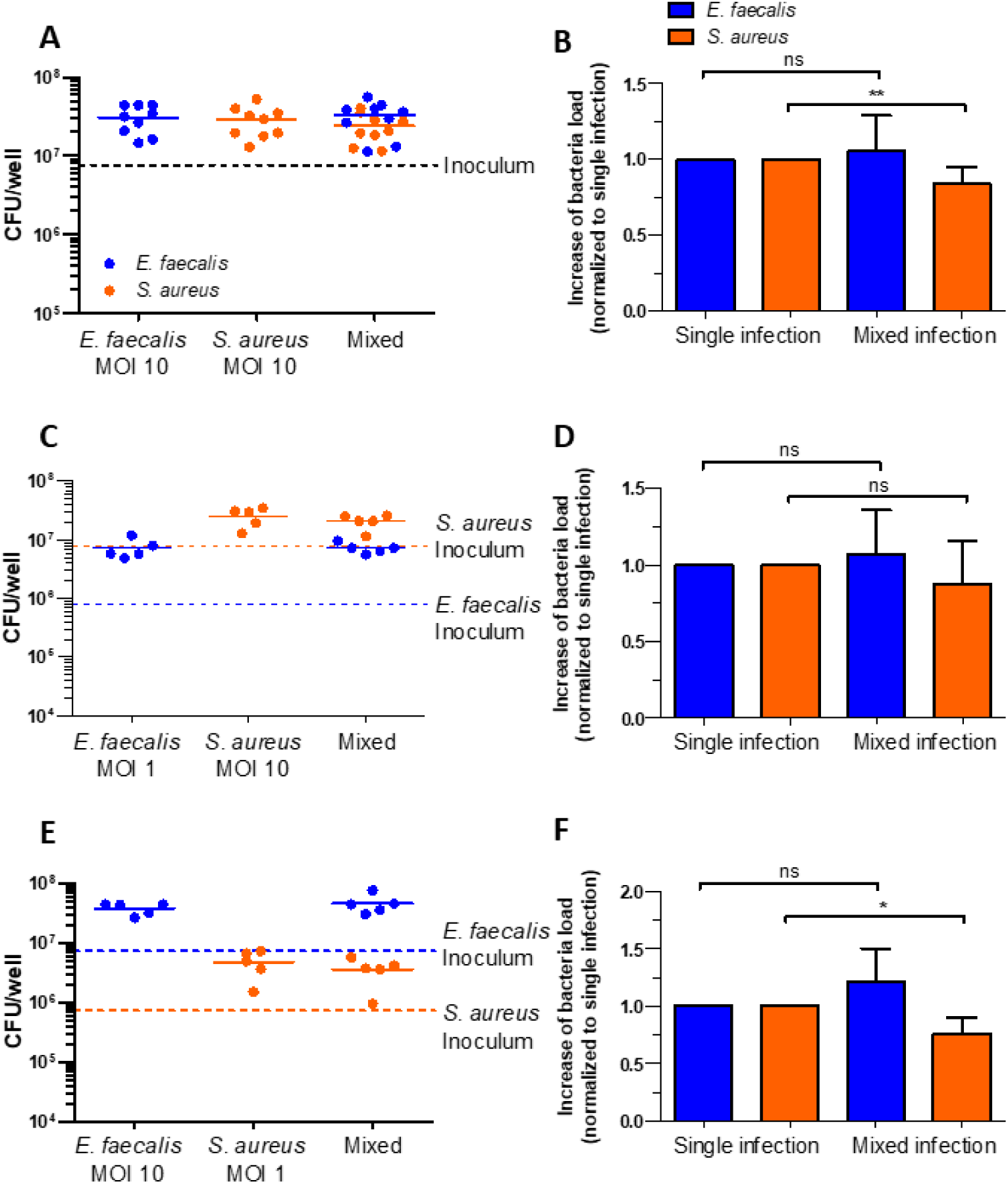
Co-culture of *E. faecalis* and *S. aureus* in the absence of neutrophils does not impact bacterial CFU. Bacterial CFU of *E. faecalis* and *S. aureus* single- and mixed-species culture of for 4-hour without neutrophils. **(A-B)** CFU (A) and fold-change (B) of *E. faecalis* (MOI 10), *S. aureus* (MOI 10), or mixed-infection (MOI 10 + 10). **(C-D)** CFU (C) and fold-change (D) of *E. faecalis* (MOI 1), *S. aureus* (MOI 10), or mixed-infection (MOI 1 + 10). **(E-F)** CFU (E) and fold-change (F) of *E. faecalis* (MOI 10), *S. aureus* (MOI 1), or mixed-infection (MOI 10 + 1). **(A**,**C**,**E)** Horizontal bars represent CFU means and dotted lines to indicate bacterial inoculum. **(B**,**D**,**F)** Bars represent means±SD for fold change relative to each single-species infection. Differences between mixed- and single-species infection were analyzed by paired t-test within each species and differences were considered significant for * p<0.05, ** p<0.01. ns = not significant.

**Supp Figure 3.**
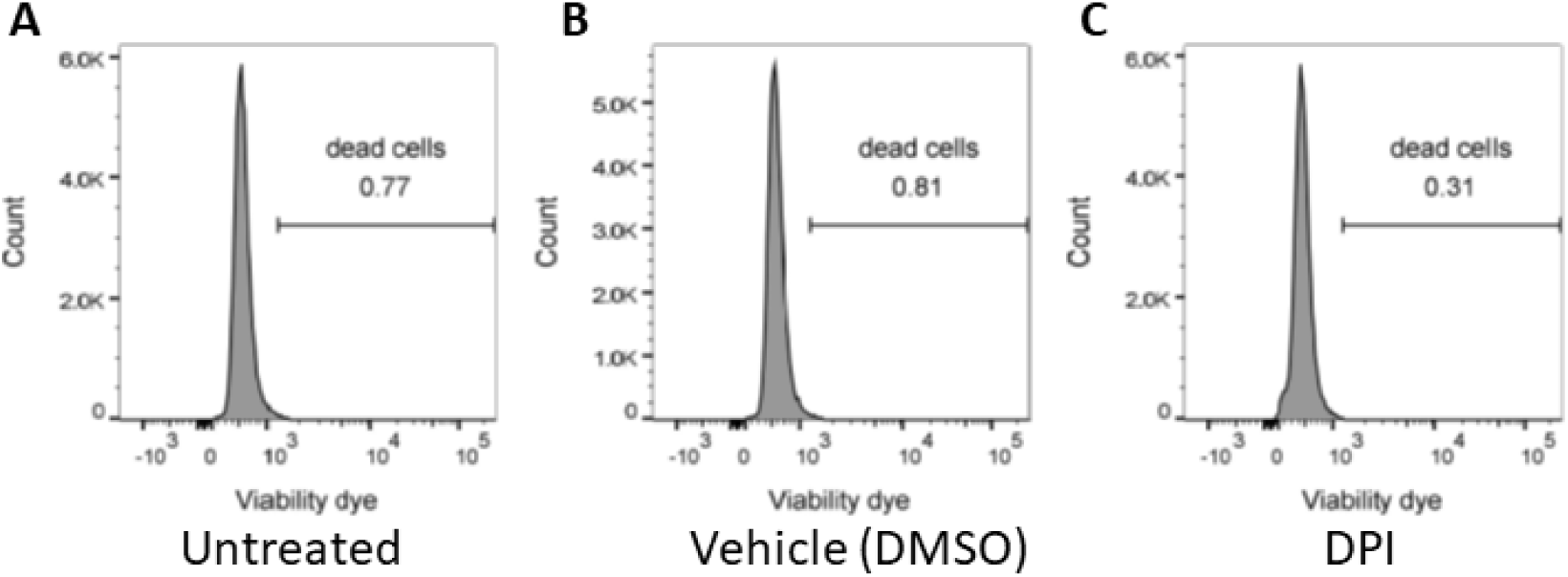
Pre-treatment of DPI does not affect neutrophil viability. Neutrophils were cultured with **(A)** no stimulation, **(B)** 0.2% of DMSO, or **(C)** 30 μM of DPI for 30 minutes at 37°C. Viability of neutrophils was measured by flow cytometry following staining with fixable viability dye from Thermofisher. Results represent the intensity of viability dye (x-axis) and cell count (y-axis). The data are the representative samples from three biological repeats.

**Supp Figure 4.**
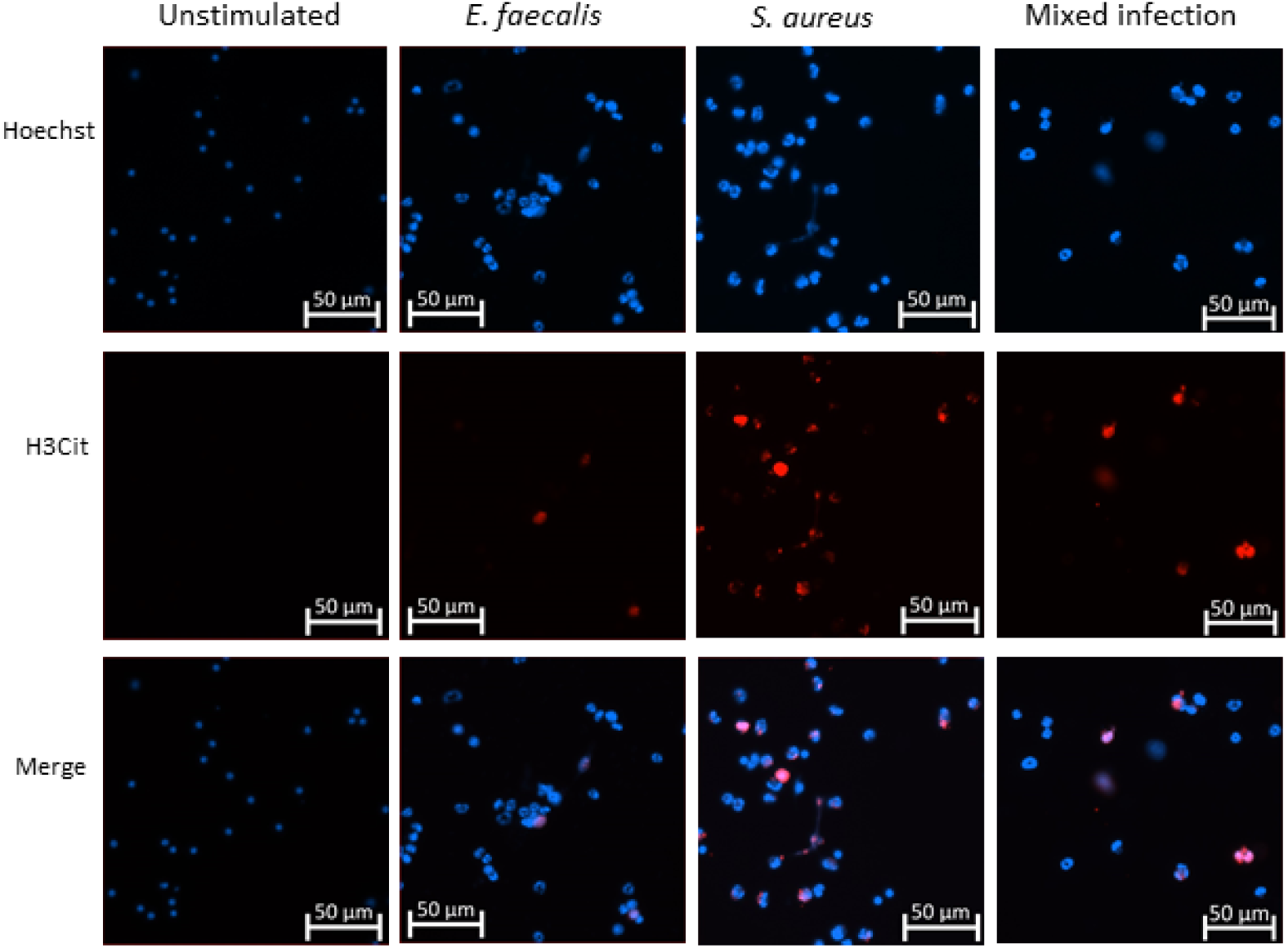
*E. faecalis* co-infection attenuates citrullination of histone H3 induced by *S. aureus*. Histone citrullination was detected by immunofluorescent staining of citrullinated histone H3 (red) and all cells were stained by Hoechst for DNA (blue). Representative images were shown with scale bar = 50μm.

**Supp Figure 5.**
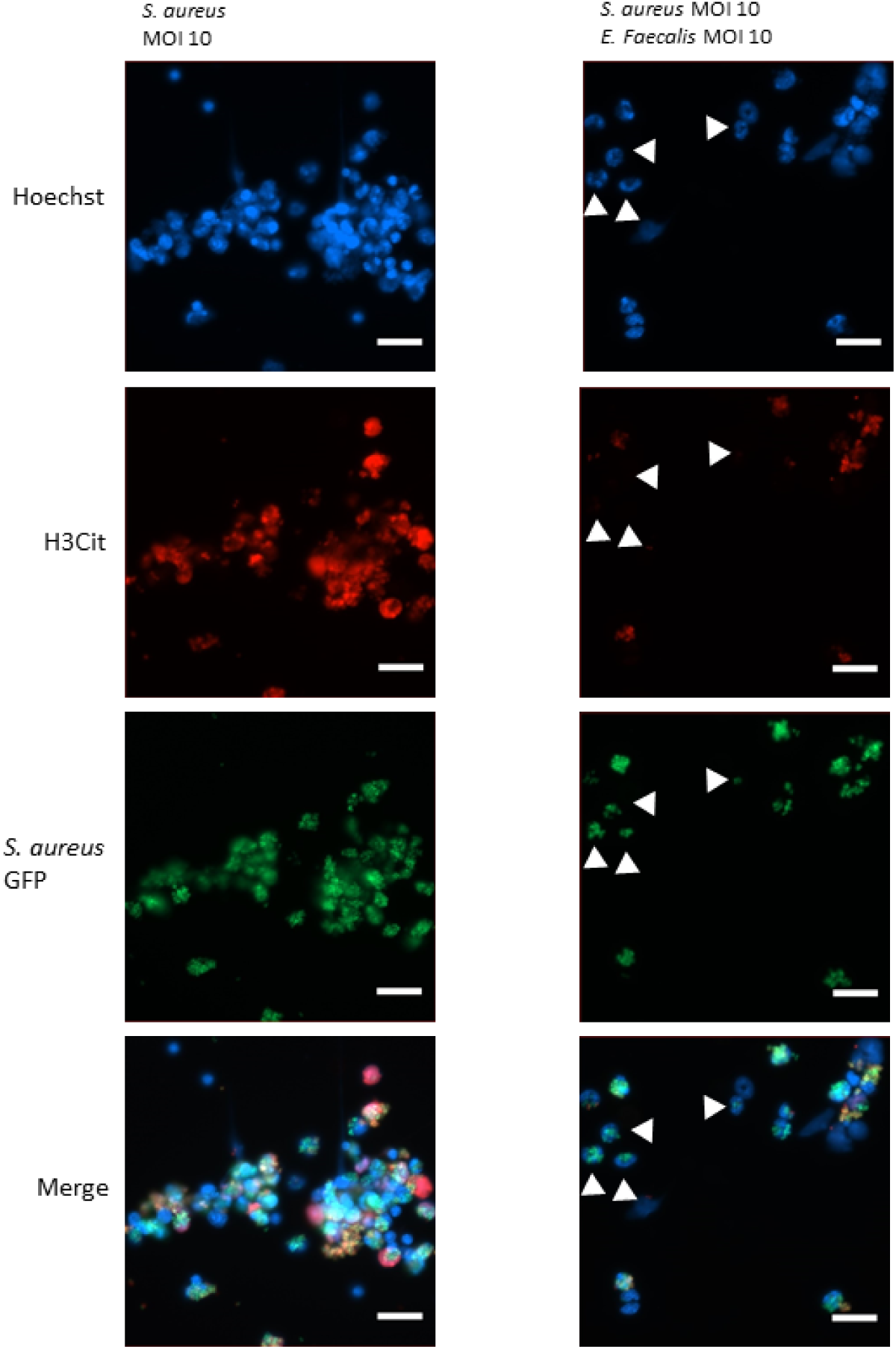
Neutrophils exhibiting citrullinated histone 3 are often associated with *S. aureus*, and co-infection with *E. faecalis* reduces the NETosis of *S. aureus*-containing neutrophils. Localization of neutrophil citrullinated histone H3 was determined by immunolabeling after 4 hours of infection with *S. aureus* in the presence or absence of *E. faecalis*. Hoechst (blue) was used to stain DNA and localization of citrullinated histone H3 (red) and *S. aureus* (green) were examined (scale bar = 20μm). White arrowheads indicate *S. aureus*-containing neutrophils without decondensed chromatin.

**Supp Figure 6.**
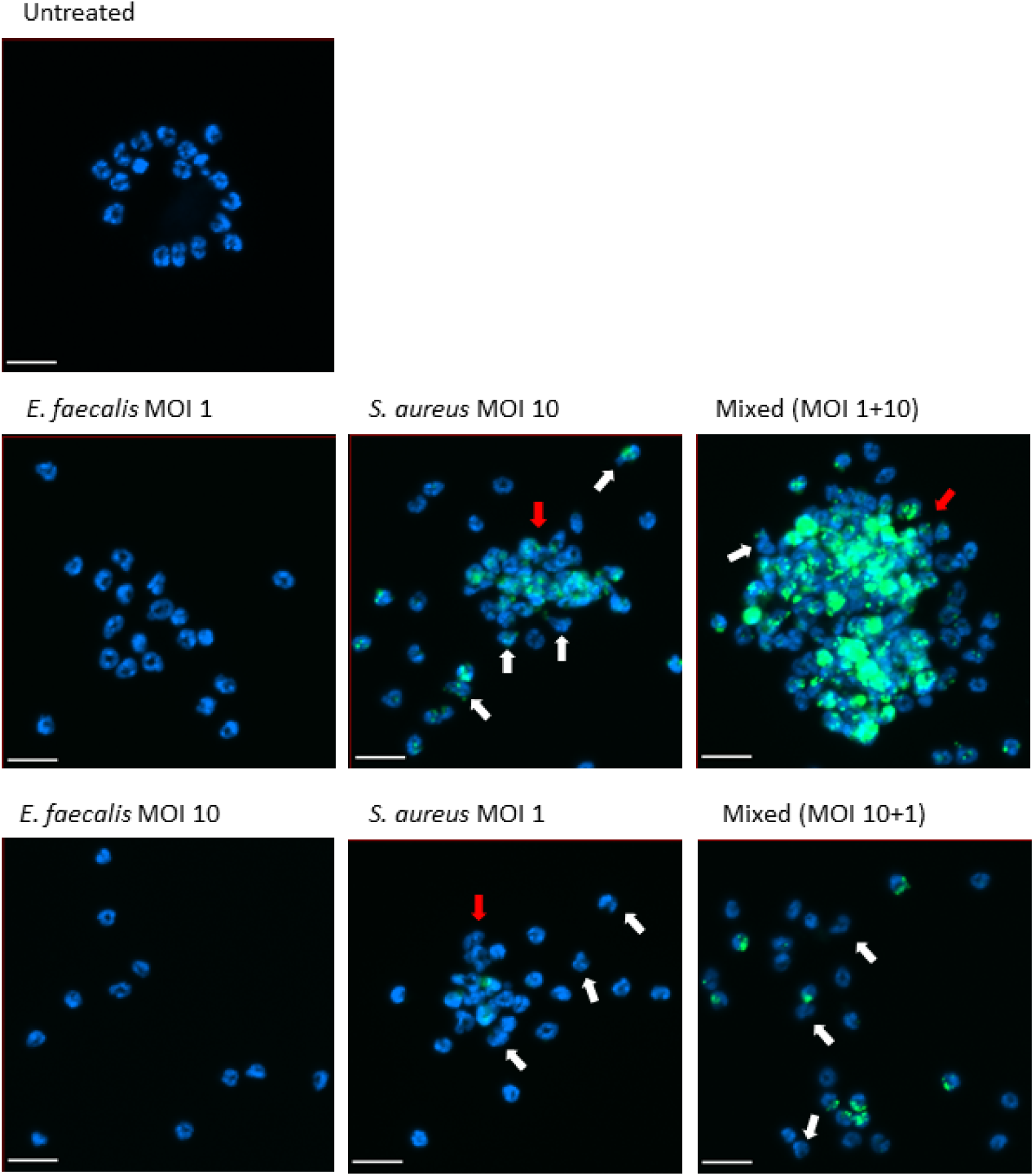
The level of NETosis in mixed-infection resembles the phenotype of single-infection by the species with the higher inoculum. Level of NETosis was measured by imaging the morphology and aggregation of neutrophils. Neutrophils were incubated with the bacteria indicated for 4 hours. Hoechst (blue) was used to stain nucleus and co-localization with GFP-expressing *S. aureus* (green) was examined (scale bar = 20μm).

**Graphical abstract.**
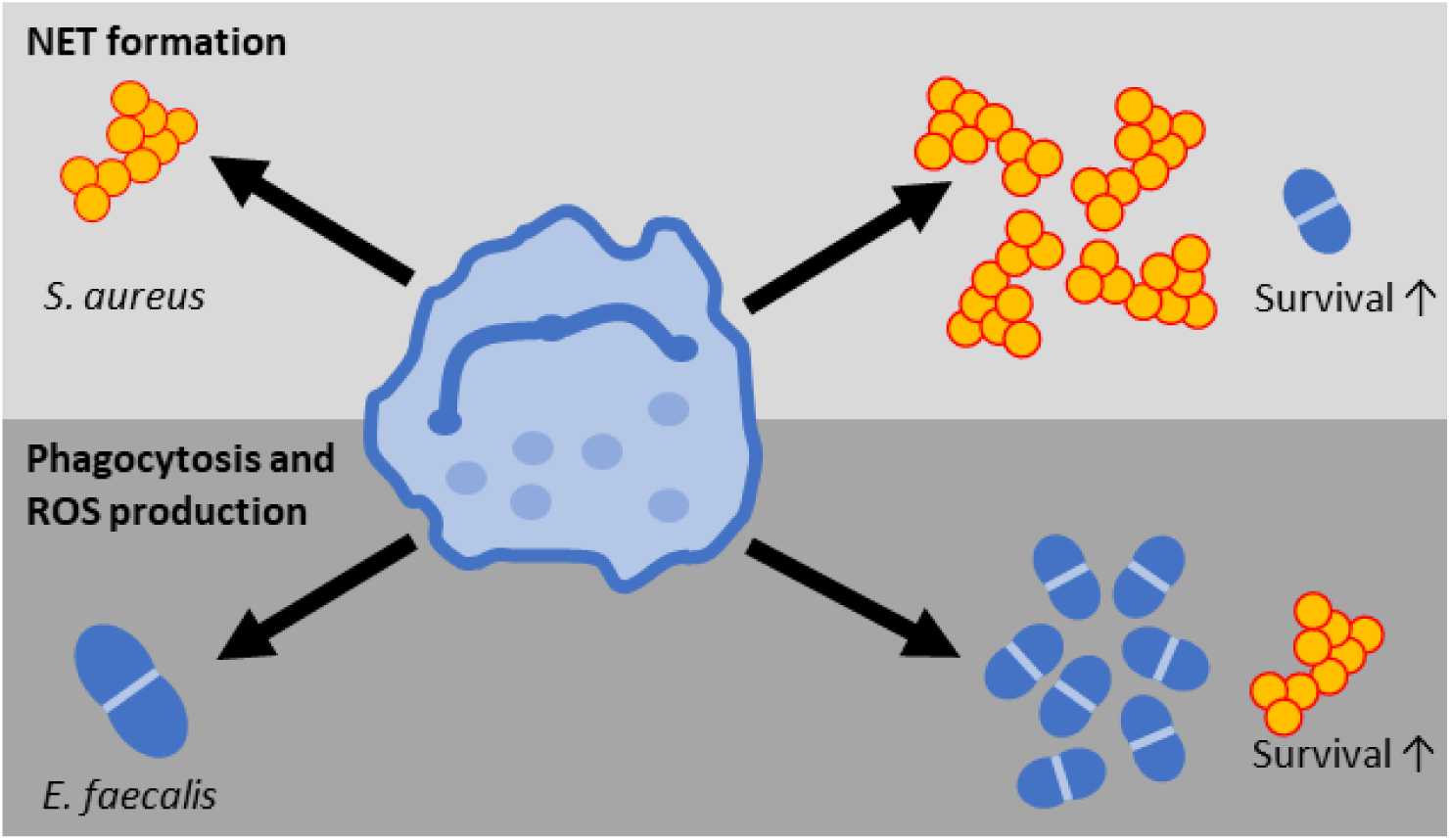
In mixed-infection, pathogen-specific antimicrobial activities are directed to respond to the predominant species and favors the survival of the less abundant species.

